## Supplementary for "Causal evidence for the adaptive benefits of social foraging in the wild"

**Supplementary Results**

(a) Likelihood of patch discovery

(b) Patch discovery latency

(c) Recruitment latency

**Supplementary Tables 1 & 2**

**Supplementary Figures 1 - 10**

**Supplementary Results:**

***Analysis at the batch level***

A necessary—but not sufficient—condition for the adaptive benefits of socially mediated resource detection is that food patches are found more often, and/or quicker, by *any* individual, the higher the number of individuals in a pool. To verify this, we ran an additional set of models at the level of the batch. These models (presented below) follow the same model structure as models I and II described in the Methods, excluding the main effect of body length and its interaction with sex-composition. Since these models were conducted on the batch level and not the individual level, individual identity as random effect was also dropped.

(a) **Likelihood of patch discovery**

As expected, the likelihood of a batch (i.e. any fish in a batch) discovering a novel food patch increased with the number of conspecifics in the pool (Estimate (*Est*) ± Standard Error (*SE*) = 1.10 ± 0.09, *N* = 1450, *χ^2^* = 89.06, *p* < 0.001; Supplementary Figure 9), independent of whether the batch consisted of males or females (Interaction effect: *Est* ± *SE* = −0.050 ± 0.174, *N* = 1450, *χ^2^* = 0.08, *p* = 0.78; Supplementary Figure 9). Male and female batches were also equally likely to discover patches overall (*Est* ± *SE* = −0.048 ± 0.168, *N* = 1450, *χ^2^* = 0.08, *p* = 0.78). There was no effect of trial number (*Est* ± *SE* = −0.01 ± 0.06, *N* = 1450, *χ^2^* = 0.02, *p* = 0.90; Supplementary Figure 10) or pool surface area (*Est* ± *SE* = 0.02 ± 0.17, *N* = 1450, *χ^2^* = 0.01, *p* = 0.92) on the likelihood of a batch discovering a novel food patch.

(b) **Patch discovery latency**

As expected, larger batches also discovered novel food patches faster (*Hazard ratio* (*95%* Confidence Interval) = 2.23 (1.98–2.51), *N*_not censored_/*N*_total_ = 599/1378, *χ2* = 92.64, *p* < 0.001; Supplementary Figure 11). However, due to the effect of fish number becoming weaker with time within a trial (0–120 seconds), the proportional hazards assumption was not met. To nevertheless evaluate the potential effect of sex-composition and trial number, we therefore stratified fish number, allowing for different baseline hazards for different fish number categories. Male and female batches did not differ in how quickly they discovered a patch (*Hazard ratio* (*CI*) = 0.94 (0.71–1.24), *N* = 599/1378, *χ2* = 0.24, *p* = 0.63; Supplementary Figure 11). Trial number did not affect how quickly a patch was discovered (*Hazard ratio* (*CI*) = 0.99 (0.91–1.07), *N* = 599/1378, *χ2* = 0.08, *p* = 0.78), neither did pool surface area (*Hazard ratio* (*CI*) = 0.94 (0.74–1.20), *N* = 599/1378, *χ2* = 0.27, *p* = 0.60). When we excluded the largest fish number category (five to eight fish), thereby meeting the proportional hazards assumption, there was still a clear positive effect of fish number (up to four fish) (*Hazard ratio* (*CI*) = 1.61 (1.40–1.84), *N* = 299/964, *χ2* = 31.35, *p* < 0.001). Again, there was no main effect of sex-composition (*Hazard ratio* (*CI*) = 0.84 (0.63–1.12), *N* = 299/964, *χ2* = 1.37, *p* = 0.24).

(c) **Recruitment latency**

To compare the recruitment latency (time between arrival of the first and second fish) of male and female compositions, we considered only patch discoveries made within the first 60 seconds (78% of the trials in which a patch was discovered). Additionally, to meet the model assumptions, we split the analysis into two models, one for relatively small batches (2–4 fish) and one for relatively large batches (5–8 fish). If a second fish did not arrive within the two-minute trial time, a maximum recruitment latency was assigned (120 seconds – arrival latency first fish) and the value was treated as censored.

Females and males did not differ in recruitment speed, either in relatively small (*Hazard ratio* (*CI*) = 0.79 (0.47–1.32), *N* = 82/118, *χ2* = 0.82, *p* = 0.37; Supplementary Figure 12) or large batches (*Hazard ratio* (*CI*) = 0.96 (0.67–1.36), *N* = 203/250, *χ2* = 0.06, *p* = 0.80; Supplementary Figure 12). Trial number and pool surface area did not affect recruitment latency in either of the models (all *p* > 0.2).

**Supplementary Table 1.** Size characteristics and number of batches per treatment type for each pool. Depth was calculated as the average depth of the five locations in each pool.

|  |  |  | **Number of fish** | | | | | | | |
| --- | --- | --- | --- | --- | --- | --- | --- | --- | --- | --- |
|  |  |  | **Females** | | | | **Males** | | | **Control** |
| **Pool** | **Surface (m^2^)** | **Depth**  **(m)** | **1** | **2–4** | **5–8** | **1** | | **2–4** | **5–8** | **5–8** |
| **P1** | 3.81 | 0.16 | 2 | 1 | 2 | 2 | | 2 | 2 | 1 |
| **P2** | 2.85 | 0.14 | 3 | 1 | 1 | 3 | | 1 | 1 | 1 |
| **P3** | 3.37 | 0.26 | 2 | 3 | 1 | 2 | | 1 | 2 | 2 |
| **P4** | 4.59 | 0.19 | 3 | 2 | 1 | 2 | | 2 | 1 | 1 |
| **P5** | 2.59 | 0.14 | 2 | 2 | 2 | 2 | | 1 | 2 | 2 |
| **P6** | 3.33 | 0.12 | 2 | 1 | 3 | 1 | | 1 | 1 | 2 |
| **P7** | 2.43 | 0.15 | 2 | 1 | 2 | 3 | | 1 | 2 | 1 |
| **Total** |  |  | 16 | 11 | 12 | 15 | | 9 | 11 | 10 |

**Supplementary Table 2.** Number of batches per treatment level (and number of focal individuals in brackets). Note that in the control treatment, we were only interested in male behavior (one per batch), explaining why the number of focal individuals equals the number of batches in this treatment.

|  | **Fish number** | | | | | | | |
| --- | --- | --- | --- | --- | --- | --- | --- | --- |
| **Composition** | **1** | **2** | **3** | **4** | **5** | **6** | **7** | **8** |
| **Female** | 16 (16) | 2 (4) | 4 (12) | 5 (20) | 2 (10) | 3 (18) | 4 (28) | 3 (24) |
| **Male** | 15 (15) | - | 5 (15) | 4 (16) | - | - | 1 (7) | 10 (80) |
| **Control**  One male + females | - | - | - | - | 1 (1) | 1 (1) | 3 (3) | 5 (5) |

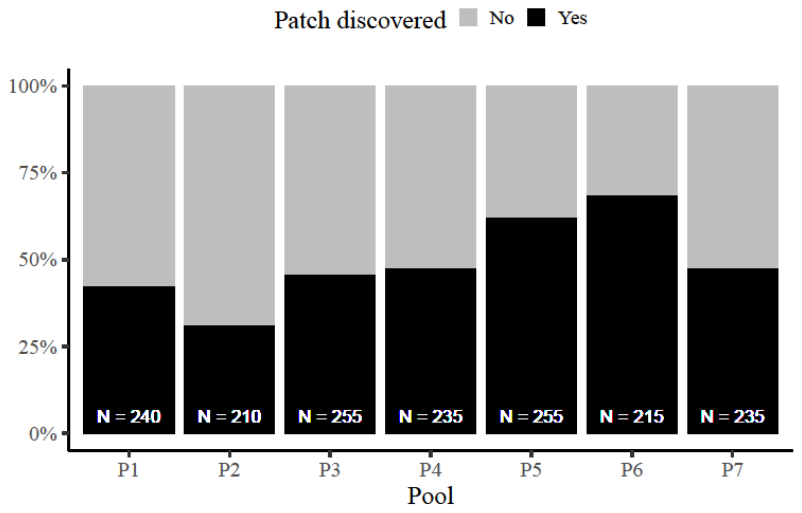

**Supplementary Figure 1.** Percentage of novel food patches discovered by a batch in each of the pools. Numbers indicate the number of trials per pool, including trials with control (i.e. mixed-sex) batches.

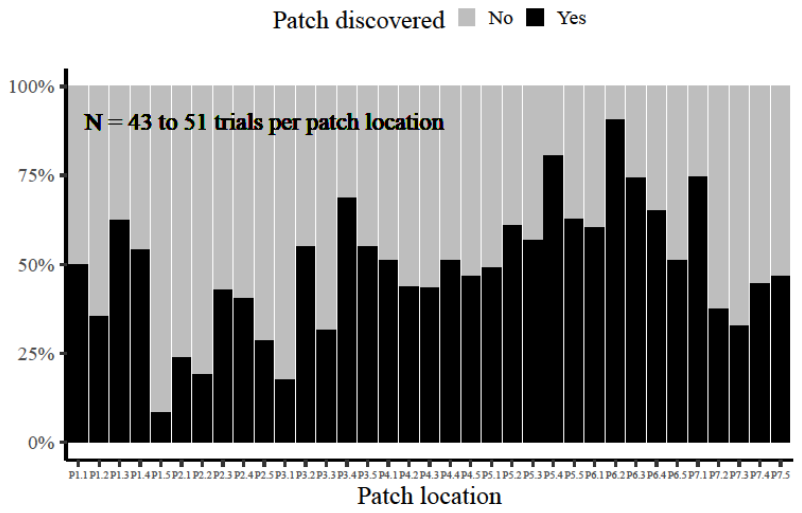

**Supplementary Figure 2.** Percentage of novel food patches discovered by a batch in each patch location (five locations per pool), including trials with control (i.e. mixed-sex) batches.

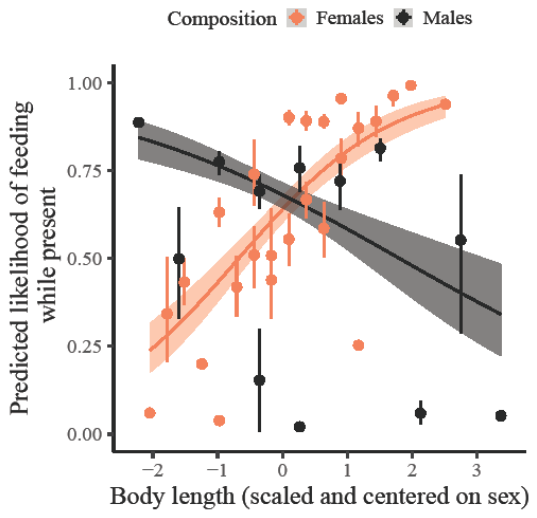

**Supplementary Figure 3.** Predicted probability of an individual feeding when at a patch as a function of its body length. Dots with bars represent the mean ± 95% confidence interval (*CI*) summary statistics for each sex and body length (*CI* obtained from 1,000 bootstraps). Grouping of the data was conducted for graphical purposes only; analyses were conducted on an individual-by-trial level with the number of fish in the pool as a continuous variable. Regression lines show the predicted final model values. Shaded areas around the lines reflect 95% *CI*. A slight horizontal position dodge was added to reduce overlap.

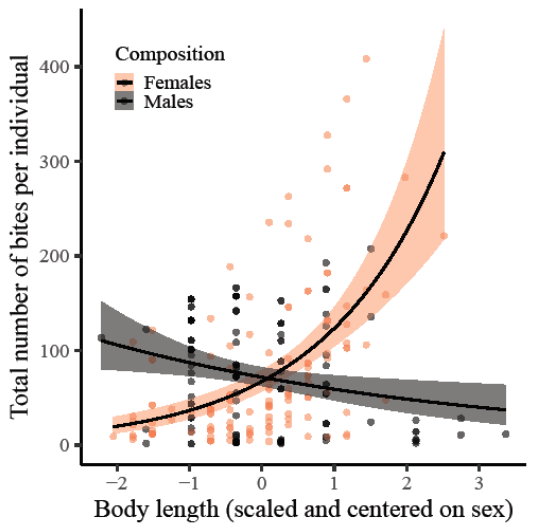

**Supplementary Figure 4.** Total number of foraging bites per individual as a function of its body length. Regression lines and 95% confidence intervals (shaded area) are based on fitted values.

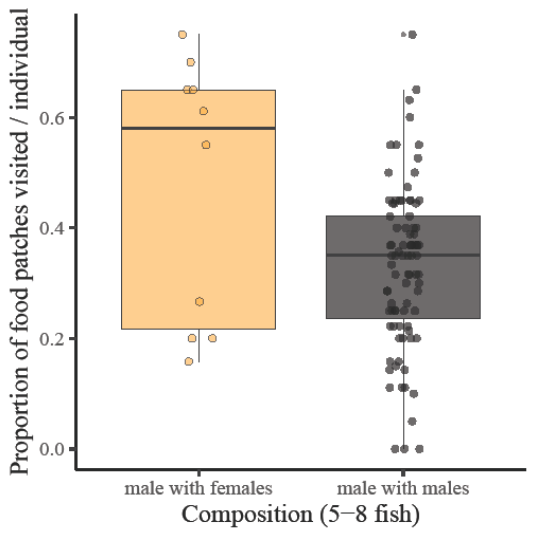

**Supplementary Figure 5.** Proportion of trials in which a novel food patch was visited by a male as a function of sex-composition. Grouping of the data on individual level (individual dots) was conducted for graphical purposes only; analyses were conducted on an individual-by-trial level. Box plots show median and interquartile range with whiskers of 1.5 interquartile distances. A slight jitter was added to the dots to reduce overlap.

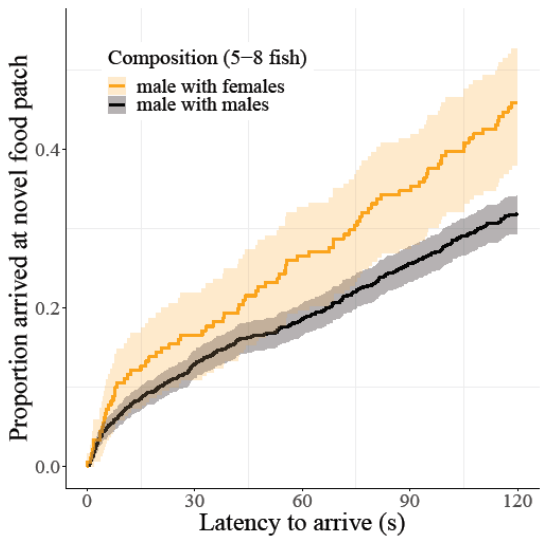

**Supplementary Figure 6.** Proportion of trials in which males arrived at a novel food patch as a function of trial time and sex composition. Higher values on the y-axis reflect an increasing number of individuals arriving at a food patch by that time. Shaded areas around the lines reflect 95% confidence intervals.

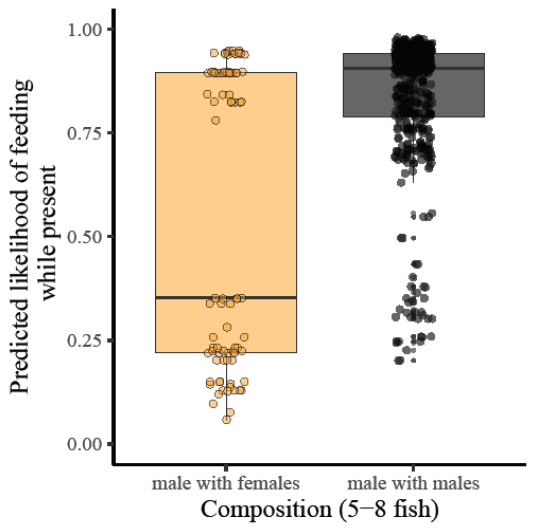

**Supplementary Figure 7.** Predicted probability of a male individual feeding at a patch as a function of the sex-composition of the batch. Analyses were conducted on an individual-by-trial level. The bimodal distribution in the mixed composition is driven by three males feeding during more than ten trials while the other seven males only fed in a maximum of three trials. Box plots show median and interquartile range with whiskers of 1.5 interquartile distances. A slight jitter was added to the dots to reduce overlap.

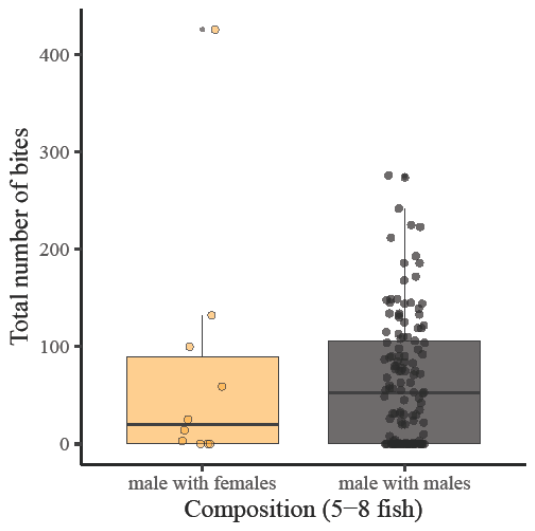

**Supplementary Figure 8.** Total number of foraging bites per male individual as a function of the sex-composition of the batch. Box plots show median and interquartile range with whiskers of 1.5 interquartile distances. A slight jitter was added to the dots to reduce overlap.

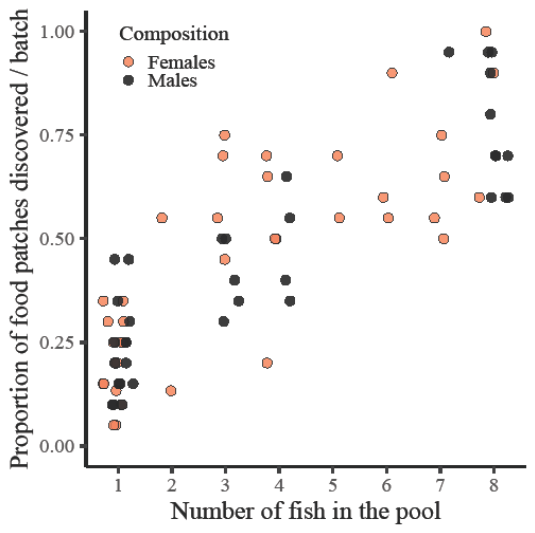

**Supplementary Figure 9.** Proportion of trials in which a novel food patch was discovered by at least one fish in the batch as a function of the number of fish in the pool. Grouping of the data on the batch level (individual dots) was conducted for graphical purposes only; analyses were conducted on a trial level. A slight jitter was added to the dots to reduce overlap.

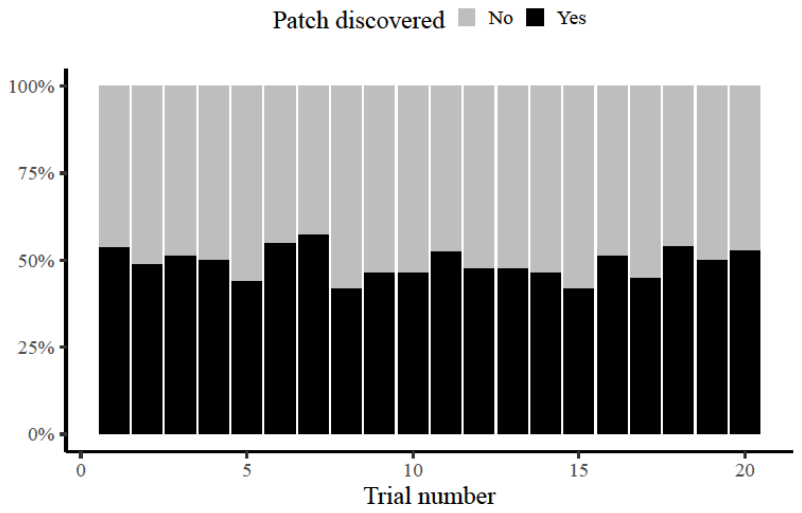

**Supplementary Figure 10.** Percentage of novel food patches discovered by a batch as a function of trial number, including trials with control (i.e. mixed-sex) batches.

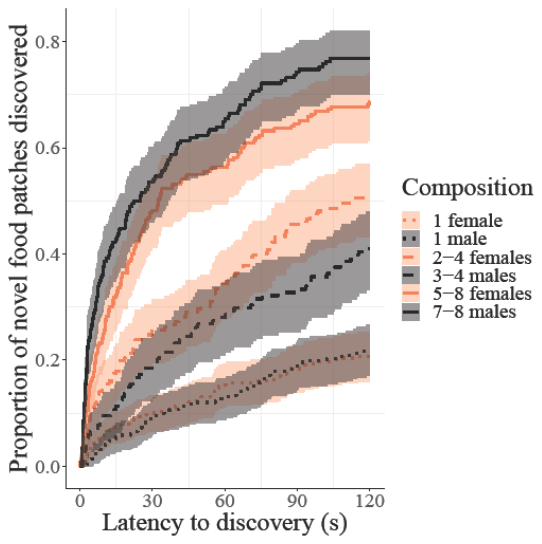

**Supplementary Figure 11.** Proportion of novel food patches discovered as a function of time in the trial for different sex-compositions and number of fish in the pool. Higher values on the y-axis reflect an increasing number of patches being discovered by that time. Grouping of the number of fish in the pool was conducted for graphical purposes only; analyses were conducted with fish number as a continuous covariate. Shaded areas around the lines reflect 95% confidence intervals.

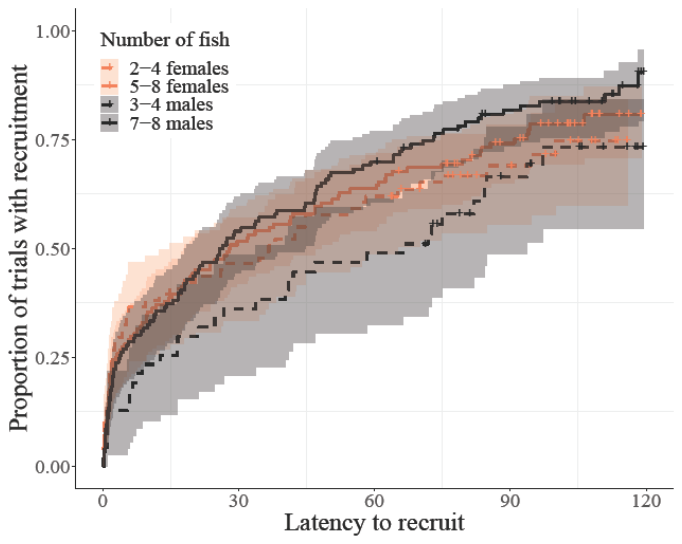

**Supplementary Figure 12.** Proportion of trials in which a second fish arrived at a discovered food patch as a function of time since its discovery. Higher values on the y-axis reflect an increasing probability of at least one individual being recruited at a discovered food patch by that time. Small plus signs on the curves indicate the presence of censored data. Shaded areas around the lines reflect 95% confidence intervals.
